# Initiation of droxidopa during hospital admission for management of refractory neurogenic orthostatic hypotension in severely ill patients

**DOI:** 10.1101/437319

**Authors:** Katherine E. McDonell, Brock A. Preheim, Andre’ Diedrich, James A.S. Muldowney, Amanda C. Peltier, David Robertson, Italo Biaggioni, Cyndya A. Shibao

## Abstract

**Introduction:** Orthostatic hypotension (OH) is a common cause of hospitalization, particularly in the elderly. Hospitalized patients with OH are often severely ill, with complex medical comorbidities and high rates of disability. Droxidopa is a norepinephrine precursor approved for the treatment of neurogenic OH (nOH) associated with autonomic failure that is commonly used in the outpatient setting, but there is currently no data regarding the safety and efficacy of droxidopa initiation in medically complex patients.

**Methods:** We performed a retrospective review of patients started on droxidopa for refractory nOH while hospitalized at Vanderbilt University Medical Center between October 2014 and May 2017. Primary outcome measures were safety, change in physician global impression of illness severity from admission to discharge, and persistence on medication after 180-day follow-up.

**Results:** A total of 20 patients were identified through chart review. Patients were medically complex with high rates of cardiovascular comorbidities and a diverse array of underlying autonomic diagnoses. Rapid titration of droxidopa was safe and well-tolerated in this cohort, with no cardiovascular events or new onset arrhythmias. Supine hypertension requiring treatment occurred in four patients. One death occurred during hospital admission due to organ failure associated with end-stage amyloidosis. Treating physicians noted improvements in presyncopal symptoms in 80% of patients. After 6 months, 13 patients (65%) continued on droxidopa therapy.

**Conclusion:** In a retrospective cohort of hospitalized, severely ill patients with refractory nOH, supervised rapid titration of droxidopa was safe and effective. Treatment persistence was high, suggesting that symptomatic benefit extended beyond acute intervention.

## Introduction

Neurogenic orthostatic hypotension (nOH) is a chronic debilitating condition that affects the elderly caused by a wide variety of neurodegenerative conditions including diabetes, cardiovascular disease, Parkinson disease (PD) and multiple system atrophy (MSA), and pure autonomic failure (PAF).

Patients affected by nOH are often medically complex with multiple comorbidities and have relatively high rates of hospital admissions, with the highest rate of 233 per 100,000 observed in adults over age 75^1^. OH is associated with an increased risk of falls^2^, decreased quality of life, and an increased rate of all-cause mortality^3^. A recent retrospective study of 1700 patients with OH from the TennCare database revealed that over 60% had been hospitalized in the previous 30 days, 50% were considered disabled, and 17% were nursing home residents^4^. High rates of cardiovascular comorbidities including hypertension (62%), syncope (40%), diabetes (38.5%), and congestive heart failure (26%) were also noted. On average, patients had used 14 different medications within the previous 6 months, reflecting the high medical complexity of this patient population.

Droxidopa is a norepinephrine precursor that recently became the second medication approved for the treatment of nOH in 2014. It has been shown to be effective for the treatment of symptomatic nOH resulting from PD, MSA, PAF, dopamine β-hydroxylase deficiency, and non-diabetic autonomic neuropathy^5–8^. The most commonly observed adverse effects in these trials included headache, dizziness, nausea, and hypertension. However, the efficacy of droxidopa has only been demonstrated over two weeks, and data on long-term safety and efficacy is still forthcoming^9^.

Droxidopa is typically initiated in the outpatient clinic through a specialty pharmacy and requires careful dose titration and monitoring. However, in a real-world setting, patients with severe nOH often have multiple comorbidities including hypertension, heart failure, diabetes mellitus, and end-stage renal disease. Droxidopa carries warnings regarding supine hypertension as well as the risk of exacerbating existing ischemic heart disease, arrhythmias, and congestive heart failure, and its safety has not been established in patients with these conditions^10^. Furthermore, given the high rates of comorbidities in patients with OH, guidance is needed regarding the safety and efficacy of rapid droxidopa initiation in the inpatient setting.

The majority of nOH patients admitted to the hospital have refractory hypotension and nOH, which are often associated with prolonged hospitalizations and severe de-conditioning. Shortly after the approval of droxidopa in October 2014, the Vanderbilt Pharmacy and Therapeutics Committee approved droxidopa for inpatient use and added the medication to our hospital formulary.

Currently there are no clinical trial data that address the relative safety of droxidopa in severely ill patients with nOH, and randomized trials are difficult to conduct in this population given the rarity of this condition and medical complexity of these patients. In the absence of these data, a retrospective study from a major referral center for the management of autonomic disorders can provide valuable information to inform health care providers regarding the management of nOH in severely ill patients in the hospital setting.

The goal of this study is therefore to determine the safety and effectiveness of droxidopa initiation in severely ill patients in the inpatient setting.

## Methods

This is a cross-sectional retrospective cohort study that included patients admitted to Vanderbilt University Medical Center between October 1, 2014 and May 31, 2017 who were treated with droxidopa for autonomic impairment. Patients were included in the cohort if they met the following criteria: (i) adults ≥ 18 years of age at time of admission, (ii) nOH refractory to midodrine and/or fludrocortisone, (iii) received at least one dose of droxidopa during hospitalization. Patients were excluded if (i) droxidopa had been initiated in an outpatient setting or (ii) droxidopa was initiated for an indication other than nOH.

Comprehensive reviews of the electronic medical record (EMR) and inpatient ordering system were performed to confirm these inclusion and exclusion criteria. Additional information on clinical characteristics, laboratory values, treatments, and outcomes was collected using a structured electronic data collection form. Collection, recording, and reporting of data was conducted securely and all necessary steps were made to ensure the privacy, health, and welfare of research subjects during and after the study. This study was approved by the Institutional Review Board of Vanderbilt University.

Primary outcome measures were safety (defined as the presence of complications including arrhythmias, severe hypertension, and cardiovascular events), change in physician global impression of severity of illness from admission to discharge, and persistence on medication for a 180-day follow-up period.

## Results

### Demographic and clinical information

A total of 46 patients were identified who were evaluated at Vanderbilt University Medical Center from October 2014–June 2017 and had an active prescription for droxidopa based on electronic inpatient pharmacy records. Twenty-six patients were excluded because treatment was initiated in the outpatient setting and/or the diagnosis of nOH was not confirmed.

Twenty patients met inclusion and exclusion criteria and were enrolled in the study. The mean age was 61 (range 38-81). Sixteen were male and four were female. Sixteen patients (80%) were white, three were African American, one was Asian, and one was Hispanic **(Table 1)**.

**Table 1.** Demographic Characteristics of study Subjects

| Table 1. Demographic Characteristics of Study Subjects |  |  |  |  |  |  |
| --- | --- | --- | --- | --- | --- | --- |
| Parameters | Amyloidosis<br>N = 6 | MSA<br>N = 5 | AAN<br>N = 3 | PAF<br>N = 2 | Other<br>N = 4 | Total<br>N = 20 |
| Age (yrs, [IQR]) | 61 (57, 63) | 64 (64, 73) | 70 (54, 73) | 65 (65, 66) | 69 (67, 74) | 65 ( 61, 71) |
| Gender (%) |  |  |  |  |  |  |
| Female | 16.7 | 20 | 33.3 | 50 | 0 | 20 |
| Race (%) |  |  |  |  |  |  |
| White | 83.3 | 80 | 100 | 50 | 75 | 80 |
| Black | 0 | 20 | 0 | 50 | 25 | 15 |
| Other | 16.7 | 0 | 0 | 0 | 0 | 5 |
| Inpt. Dur.(d, ([IQR]) | 12.5 (3.8, 19.0) | 10.0 (4.0, 12.0) | 7.0 (6.5, 36.5) | 21.5 (14.3, 28.8) | 5.5 (1.8, 9.8) | 8.0 (3.3, 18.8) |
| Med # (med, [IQR]) | 13.5 (13.0, 14.8) | 15.0 (14.0, 16.0) | 14.0 (10.5, 18.0) | 10.0 (7.5, 12.5) | 25.5 (20.8, 27.0) | 14.5 (12.3, 19.0) |
| MSA, Multiple System Atrophy; AAN, Autoimmune Autonomic Neuropathy; PAF, Pure Autonomic Failure; |  |  |  |  |  |  |

All patients met criteria for OH with a documented decrease of ≥ 20mm Hg in systolic or ≥ 10mm Hg in diastolic BP upon standing for 1-3 min. Patients reported symptoms of OH including dizziness, lightheadedness, or feeling faint (n=17), syncope (n=17), falls (n=6), weakness (n=4), and significant fatigue (n=2).

Underlying causes of autonomic failure in this cohort included amyloidosis (n=6), multiple system atrophy (n=5), autoimmune autonomic neuropathy (n=3), pure autonomic failure (n=2), and others (diabetic autonomic failure [n=2], autonomic neuropathy related to end-stage renal disease [n=1], and Parkinson’s disease [n=1]) **(Table 1)**.

Cardiovascular comorbidities were common in these patients, including syncope (n=17), hypertension (n=8), congestive heart failure (n=5), atrial fibrillation (n=2), and atrial flutter (1). Two patients had had heart transplants and one was on hemodialysis. Five patients had a prior diagnosis of diabetes. Fifteen patients reported a history of autonomic neuropathy; five of peripheral neuropathy, and one had a history of dementia **(Table 2).**Other common medical comorbidities included kidney disease, cancer, and thyroid disease **(Figure 1)**.

**Table 2.** Co-morbidities of Study Subjects

| Parameters | Amyloidosis<br>N = 6 | MSA<br>N = 5 | AAN<br>N = 3 | PAF<br>N = 2 | Other<br>N = 4 | Total<br>N = 20 |
| --- | --- | --- | --- | --- | --- | --- |
| *Cardiovascular Conditions* |  |  |  |  |  |  |
| Cong. Heart Failure | 66.7 | 0 | 0 | 50 | 50 | 35% |
| Hypertension | 0 | 60 | 66.7 | 50 | 75 | 45% |
| Atrial Fib/Flutter | 16.7 | 0 | 33.3 | 0 | 25 | 15% |
| *Endocrine Conditions* |  |  |  |  |  |  |
| Diabetes | 33.3 | 0 | 33.3 | 0 | 75 | 30% |
| Fall Fractures | 0 | 0 | 0 | 0 | 50 | 10% |
| Thyroid Disease | 0 | 20 | 66.7 | 50 | 25 | 25% |
| *Neurological Conditions* |  |  |  |  |  |  |
| Parkinson's Disease | 0 | 20 | 0 | 0 | 25 | 10% |
| Peripheral Neuropathy | 66.7 | 0 | 33.3 | 50 | 25 | 40% |
| Syncope | 83.3 | 100 | 66.7 | 100 | 75 | 85% |
| *Other Chronic Conditions* |  |  |  |  |  |  |
| Cancer | 16.7 | 20 | 66.7 | 0 | 50 | 30% |
| ESRD/Kidney Disease | 33.3 | 40 | 0 | 50 | 25 | 30% |
| Transplant | 16.7 | 0 | 0 | 0 | 25 | 10% |

MSA, Multiple System Atrophy; AAN, Autoimmune Autonomic Neuropathy; PAF, Pure Autonomic Failure;

**Figure 1.**
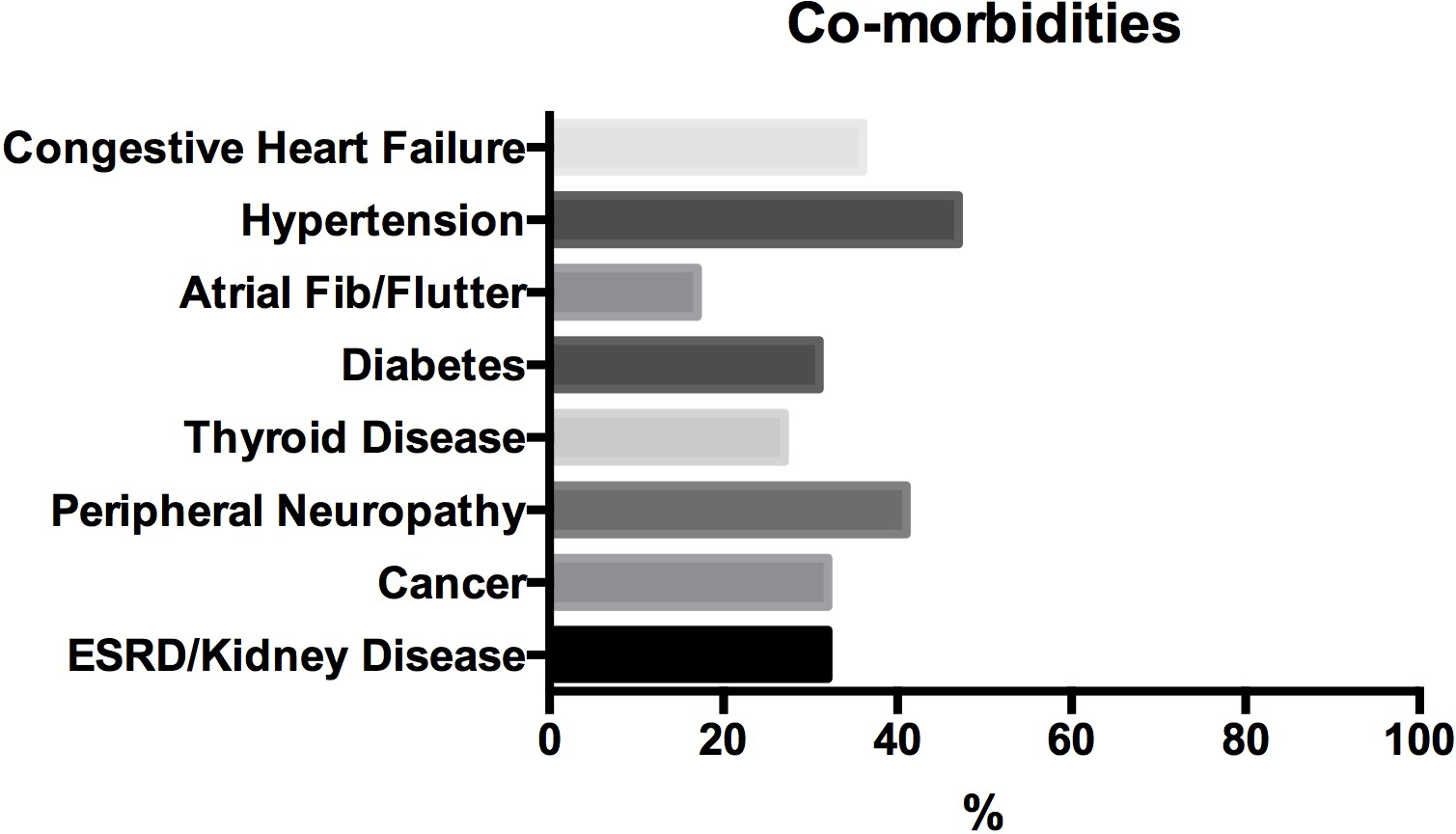
Medical comorbidities.

The median duration of inpatient hospitalization was 8 days (IQR 3-19). Median number of medications used in the past 6 months was 15 (IQR 12-19).

### Cardiovascular Autonomic Evaluation

We obtained hemodynamic data through review of the electronic database flow charts for 18 of the 20 patients. The mean baseline systolic blood pressure of the patient cohort before treatment was 129+/−30 mmHg supine and 82+/−23 mmHg after 1 minute of standing (**Figure 2**).

**Figure 2.**
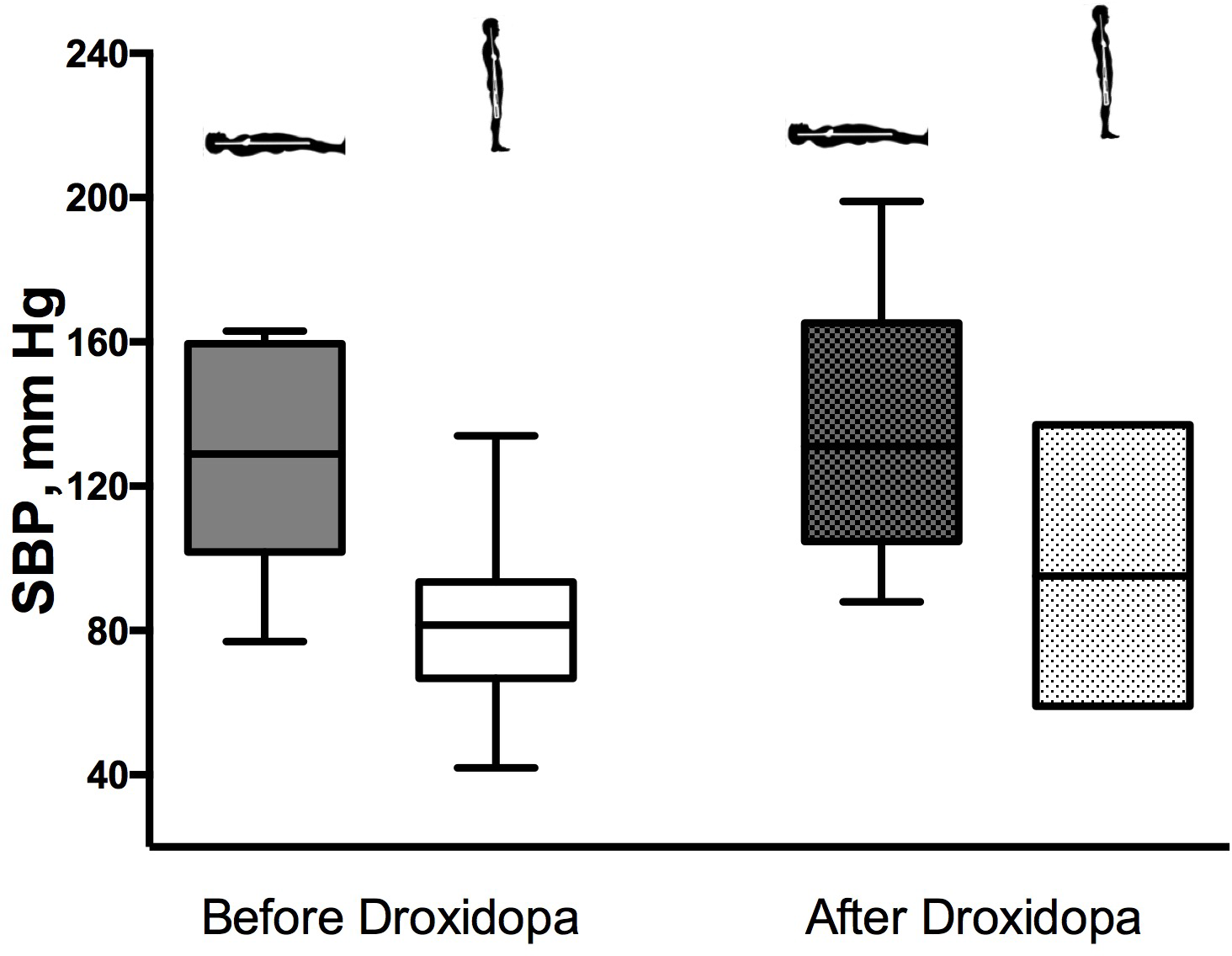
Orthostatic vital signs before and after droxidopa treatment.

### Concomitant medications

The median number of concomitant medications taken during the allotted study period was 14.5 (IQR 12.8-17.0). Of these concomitant medications, 18 out of the 20 patients in the cohort had been prescribed midodrine previously or were taking midodrine at the time of hospital admittance. 13 of the 20 had taken fludrocortisone previously or at the time of admission. 9 of the patients underwent a midodrine challenge prior to droxidopa initiation. The mean systolic blood pressure prior to midodrine treatment was 85+/−17 mmHg, and the mean systolic post-midodrine was 97+/−26 mmHg. Refractory nOH was defined as no change in standing blood pressure at 1 minute after midodrine administration^11^.

### Droxidopa initiation

All patient underwent a rapid dose titration of droxidopa over a period of 1–6 days. The average daily dose of droxidopa reached after titration was 837+/−509 mg.

### Safety

Four patients developed supine hypertension (defined as ≥150/90mmHg) requiring initiation of an anti-hypertensive medication at bedtime. One patient developed severe hypertension to 199/120, and one patient discontinued droxidopa due to hypertension.

No cardiovascular events or new onset arrhythmias were observed during rapid dose titration of droxidopa or within a two-week follow up period.

There was one death in this cohort during hospital admission. Autopsy data was consistent with organ failure due to end-stage amyloidosis.

No other adverse effects were reported in relationship to droxidopa initiation or titration.

### Clinical efficacy of droxidopa treatment

The mean systolic blood pressure at the conclusion of droxidopa titration was 138+/−31 mmHg supine and 98+/−17.5 mmHg standing for 1 minute (**Figure 2**).

We also categorized the patients by autonomic diagnosis and analyzed the hemodynamic data of these groups to determine if droxidopa is associated with a differential response in these diagnostic groups. These results are shown in **Table 3**. Overall, the amyloidosis group showed the most significant decrease in SBP drop between supine and standing on treatment (28mmHg at baseline vs. 11.2mmHg post-treatment). Patients with amyloidosis also required higher doses of droxidopa as compared with all other groups (mean daily dose 1000mg+/−577 in amyloidosis group; mean daily dose 793mg+/−460 in all others), as shown in **Figure 3**.

**Table 3.** Hemodynamic response by autonomic diagnosis

| Table 3. Hemodynamic response by autonomic diagnosis |  |  |  |  |
| --- | --- | --- | --- | --- |
| Autonomic diagnosis | Mean Baseline Supine SBP | Mean Baseline Standing SBP | Mean Post-treatment Supine SBP | Mean Post-treatment Standing SBP |
| Amyloidosis | 110+/-32 | 82+/-27 | 109.5+/-24.6 | 98.3+/-34.7 |
| MSA | 139.6+/-26 | 78.6+/-17.5 | 151.4+/-32.6 | 93.2+/-18.1 |
| AAN | 131+/-32.5 | 86+/-19.2 | 150+/-25.2 | 111+/-2.6 |
| PAF | 161.5+/-1.0 | 97+/-34 | 161+/-7.1 | 92+/-7.1 |
| Other | 123+/-2.1 | 81.3+/-14 | 133.5+/-32.1 | 94.5+/-6.9 |

MSA, Multiple System Atrophy; AAN, Autoimmune Autonomic Neuropathy; PAF, Pure Autonomic Failure.

**Table 4.** Other Pressor Medication Use

| Parameters | Amyloidosis<br>N = 6 | MSA<br>N = 5 | AAN<br>N = 3 | PAF<br>N = 2 | Other<br>N = 4 | Total<br>N = 20 |
| --- | --- | --- | --- | --- | --- | --- |
| <u>Midodrine Use</u> | 83.3 | 100 | 100 | 50 | 100 | 90 |
| Fludrocortisone Use | 33.3 | 80 | 66.7 | 100 | 75 | 65 |

MSA, Multiple System Atrophy; AAN, Autoimmune Autonomic Neuropathy; PAF, Pure Autonomic Failure.

**Figure 3.**
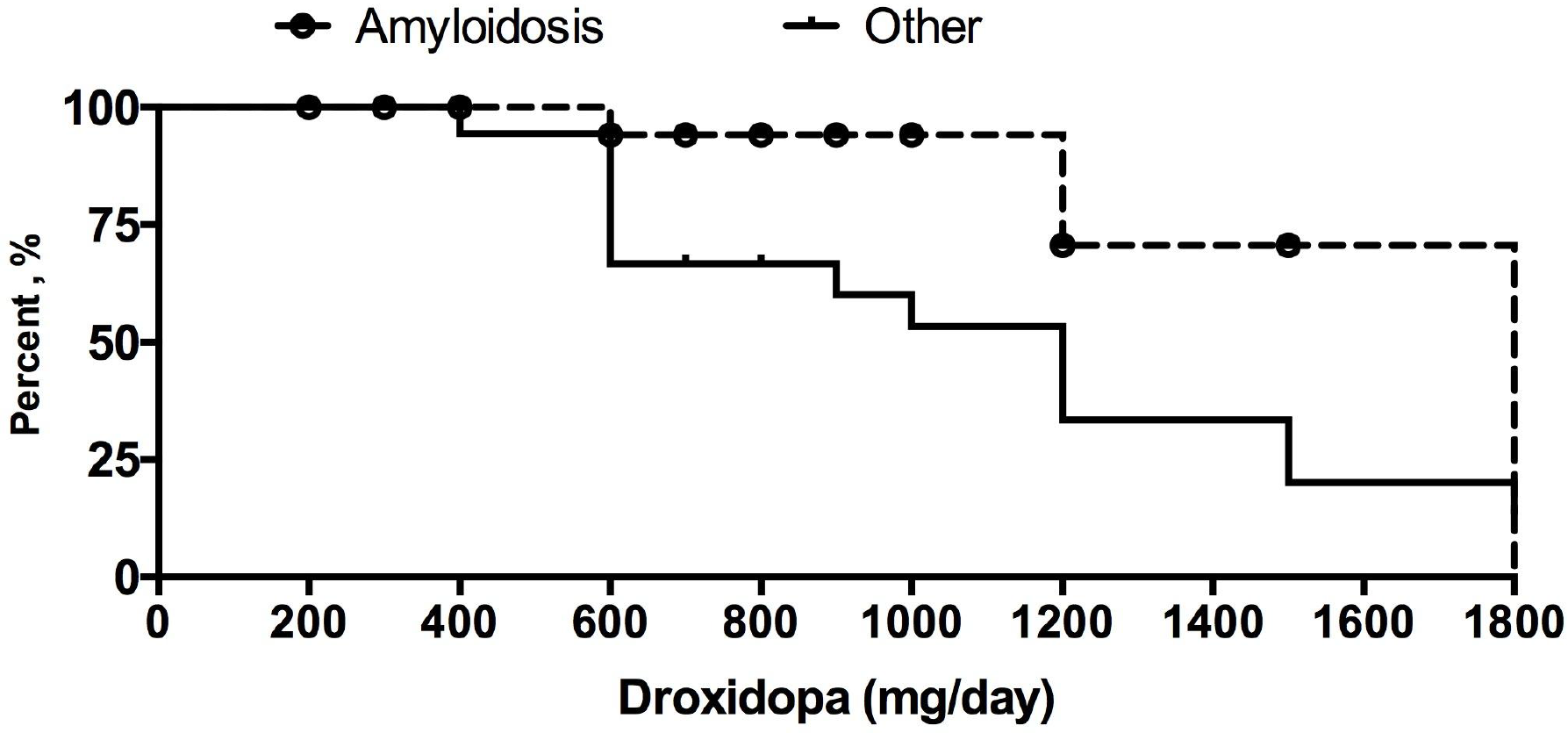
Patients with amyloidosis required higher doses of droxidopa.

### Physician judgment of response

Autonomic specialists were asked to judge the overall severity of presyncopal symptoms at admission and discharge and to rank the change in these symptoms in one of five categories: better and improved, moderately better, somewhat better, hardly any change, no change. Overall, the treating physician noted improvement in 80% of patients treated with droxidopa from admission to discharge. Significant improvement was noted in 30% of patients, somewhat to moderate improvement in 50%, and no change or hardly any change in 20% **(Figure 4)**.

**Figure 4.**
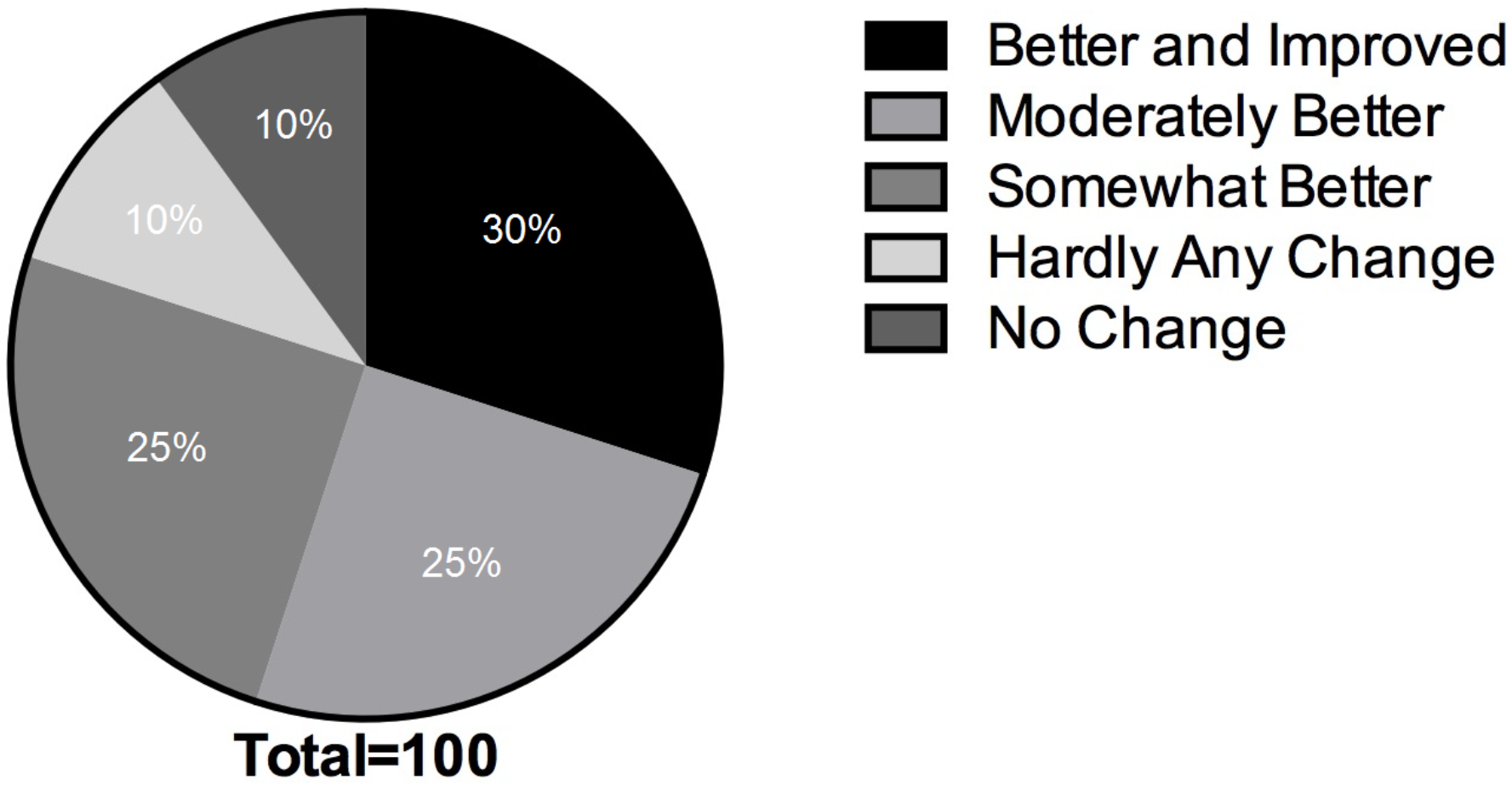
Physician judgment of response.

### Persistence on treatment

Based on follow-up chart review for 18 of the patients, 13 (65% of the initial cohort) continued droxidopa treatments after hospital discharge for >180 days **(Figure 5).**The mean persistence on treatment was 140+/−68 days. Of the five patients who discontinued droxidopa treatment prior to 180 days, three discontinued droxidopa due to side effects, one because nOH symptoms improved, and one died in inpatient care due to progression of amyloidosis. Follow up data was not available for two out of the initial twenty patients.

**Figure 5.**
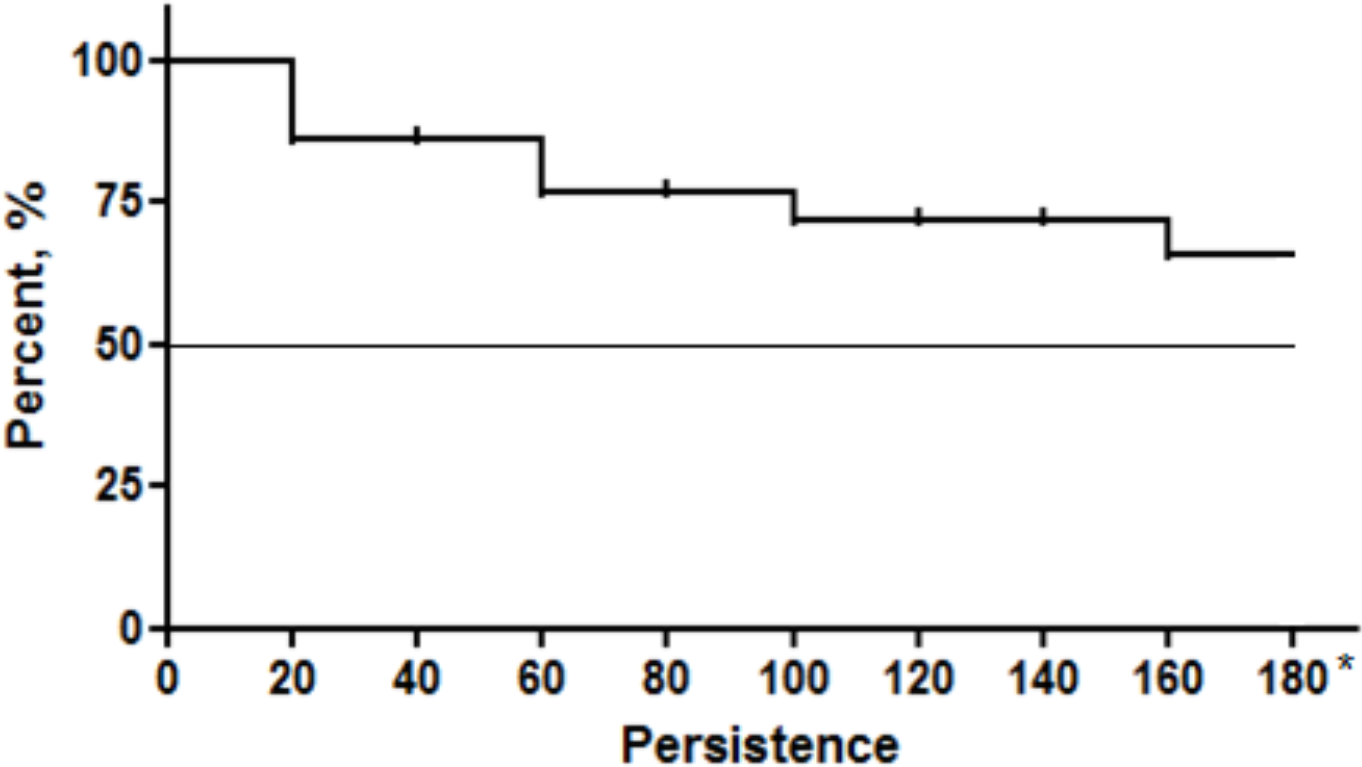
Persistence on treatment after 6 months.

## Discussion

In this retrospective cohort of hospitalized, severely ill patients with refractory nOH, droxidopa initiation was safe and was associated with clinical improvement in 80% of the treated patients. No cardiovascular events or new arrhythmias were reported during rapid dose titration. Four patients developed supine hypertension, but only one had to discontinue treatment due to this. Overall, post-treatment supine blood pressures were not significantly different from baseline (**Table 3**), but clinical improvement was still observed. Treatment persistence was also high; with 65% of patients remaining on droxidopa therapy after a 6 month follow up period.

These findings are particularly relevant given the unique and challenging nature of this population. All subjects in this study had severe, refractory nOH and multiple medical comorbidities, including significant cardiovascular comorbidities such as hypertension, congestive heart failure, and atrial fibrillation. Two had a history of heart transplant. The enrolled patients were on an average of 14 concomitant medications during their hospitalization. Based on previous data from a review the Tennessee Medicaid database, patients with nOH are a medically complex population with extremely high rates of comorbidities that make treatment challenging^4^. However, the majority of clinical trial data available is based on medically stable outpatients, providing very little guidance in terms of the management of this condition in complex, severely ill patients. Our population is likely a more representative sample that provides insight into the real world experience of treating nOH.

Data regarding the use of droxidopa in a medically complex population is particularly lacking. This medication is relatively new and is currently only the second medication approved for the treatment of nOH, along with midodrine. The long term clinical efficacy of midodrine has not been established, leading to a proposed withdrawal by the FDA in 2010 which was subsequently reversed^12^. However, in clinical practice many patients with nOH are refractory to midodrine, as was the case in our population. Additionally, the use of midodrine is relatively contraindicated in heart failure as it results in peripheral vasoconstriction, leading to increased afterload and potentially reduced systolic function. Droxidopa could therefore represent a promising alternative, given its inotropic properties. However, its safety in this population has not yet been established, and its labeling contains a warning regarding its use in patients with existing ischemic heart disease, arrhythmias, and congestive heart failure. This study therefore provides valuable data to support the use of droxidopa in a population of patients with a variety of cardiovascular comorbidities.

Our findings in the amyloidosis subgroup also warrant mention. Amyloidosis is frequently associated with severe, multifactorial orthostatic hypotension due to a combination of mechanisms, including cardiac involvement, peripheral neuropathy, and volume depletion^13^. These patients often suffer from baseline hypotension in addition to orthostasis and can be quite difficult to treat. Four out of six amyloidosis patients in our study cohort were refractory to midodrine (data was not available on the remaining two), but the majority responded positively to droxidopa, and five out of six remained on treatment after 6 months. Of note, patients with amyloidosis required higher doses of droxidopa than the rest of the cohort but showed the most significant reduction in SBP drop between supine and standing post-treatment. These findings warrant further investigation but suggest that high dose droxidopa may have a positive effect in this particularly difficult patient population.

Limitations of this study include its retrospective design and small sample size. However, even within this small sample we were able to capture a diverse array of underlying autonomic diagnoses and a high prevalence of medical comorbidities, including significant cardiovascular risk factors.

Given the current lack of clinical data regarding the use of droxidopa in these vulnerable patient populations, this study provides valuable insights to help guide clinical practice. Further longitudinal studies are needed to examine the safety and efficacy of droxidopa initiation in the inpatient setting, with longer term follow up to determine the persistence of these effects.

